# Genetic coordination of sperm morphology and seminal fluid proteins promotes male reproductive success in *Drosophila melanogaster*

**DOI:** 10.1101/2021.11.15.468624

**Authors:** Jake Galvin, Erica L. Larson, Sevan Yedigarian, Mohammad Rahman, Kirill Borziak, Michael DeNieu, Mollie K. Manier

**Author notes:** Corresponding Author: Mollie K. Manier, **Email:**. **Author Contributions:** M.K.M. designed research; J.G., E.L.L., S.Y., M.R., and M.K.M. performed research; J.G., E.L.L., K.B., M.D., and M.K.M. analyzed data; and J.G., E.L.L., and M.K.M. wrote the paper.

## Abstract

Spermatozoal morphology is highly variable both among and within species and in ways that can significantly impact fertilization success. In *Drosophila melanogaster*, paternity success depends on sperm length of both competing males and length of the female’s primary sperm storage organ. We found that genes upregulated in long sperm testes are enriched for lncRNAs and seminal fluid proteins (Sfps). Transferred in seminal fluid to the female during mating, Sfps are secreted by the male accessory glands (AG) and affect female remating rate, physiology, and behavior with concomitant advantages for male reproductive success. Despite being upregulated in long sperm testes, they have no known function in testis tissue. We found that Sex Peptide and ovulin (Acp26Aa) knockouts resulted in shorter sperm, suggesting that Sfps may regulate sperm length during spermatogenesis. However, knockout of AG function did not affect sperm length, suggesting that AG expression has no influence on spermatogenic processes. We also found that long sperm males are better able to delay female remating, suggesting higher Sfp expression in AG. These results might suggest that long sperm males have a double advantage in sperm competition by both delaying female remating, likely through transfer of more Sfps, and by resisting sperm displacement. However, we also found that this extra advantage does not necessarily translate to more progeny or higher paternity success. Thus, we found that multiple components of the ejaculate coordinate to promote male reproductive success at different stages of reproduction, but the realized fitness advantages in sperm competition are uncertain.

**Significance Statement:** The ejaculate is comprised of sperm produced in the testis and seminal fluid primarily produced in the male accessory glands (AG). These complementary components are both critical for male reproductive success, but they are largely considered to be functionally, genetically, and developmentally independent. In a quest to understand genetic mechanisms of sperm length variation, we found that genes upregulated in long sperm testes are enriched for lncRNAs and seminal fluid proteins (Sfps). Knockout of two Sfps, Sex Peptide and ovulin, results in shorter sperm, though knockout of AG function has no effect. Moreover, long sperm males delay female remating longer. These results suggest sophisticated testis-AG coordination that amplifies male reproductive success, with implications for evolutionary integration of sexually selected traits.

## Introduction

Understanding how diversity arises and is maintained is a central goal of evolutionary biology. Spermatozoa are among the most diverse cell types and have been the focus of many studies seeking to understand selective principles driving their evolution. The most familiar sperm bauplan typically features a head, a midpiece housing the mitochondria, and a flagellum tail, but variations include up to 100 flagella, no flagella, helical heads (1, 2), undulating membranes, radial symmetry, amoeboid motility (3), immobility, multiple sperm morphs from a single male, and conjugated multi-sperm structures thought to behave cooperatively (4, 5). Evolutionary forces driving such extreme diversification remain poorly understood but are thought to be related to factors like fertilization mode (6), the fertilization environment mediated by the female (7–12), and postcopulatory sexual selection (13–17). A full understanding of sperm evolutionary diversification is impossible without understanding its development, yet we know relatively little about how regulatory divergence in spermatogenesis contributes to sperm phenotypic diversity.

In *Drosophila* fruit flies, sperm length varies over two orders of magnitude from 224 μm in *D. subobscura* (18) to 58,290 μm in *D. bifurca* (19). Within *D. melanogaster*, sperm length is under postcopulatory sexual selection, with complex interactions mediating the outcome of both sperm competition and cryptic female choice. Sperm length interacts with sperm numbers as well as with length of the primary female sperm storage organ, the seminal receptacle (SR), in a way that is contingent on phenotypes of the first male, second male, and female (7, 12, 20–22). Specifically, the effect of sperm length on fertilization success depends on SR length, such that longer sperm have a competitive advantage in long SRs, while shorter sperm are advantageous in short SRs (12, 20). Sperm length and SR length are positively correlated across *Drosophila* species (23), a pattern that may be mediated by these functional sperm-SR interactions as well as by a genetic correlation between the two traits (22). In terms of other fitness effects, both long sperm and long SRs are associated with enhanced longevity and few overall fitness costs (24). However, trade-offs and condition-dependence of sperm length become more apparent in species with extremely long sperm, consistent with giant sperm evolving as an exaggerated sexual ornament (22). Indeed, runaway selection may be an important factor in sperm length evolution, fueled by the genetic correlation between the female choice trait (SR length) and the male ornament (sperm length) (22).

A key missing component in our understanding of sperm length evolution is knowing how sperm elongation is developmentally regulated during spermatogenesis. In *D. melanogaster*, spermatogenesis begins at the apical tip of the testis, when progenitor stem cells undergo asymmetrical mitosis to yield a diploid spermatogonium. This spermatogonium is born enclosed within two somatic cyst cells that all together comprise the cyst, the primary unit of synchronous spermatogenesis. The spermatogonium completes four rounds of mitosis, yielding 16 spermatocytes that undergo a period of dramatic growth and transcription, followed by meiosis to yield 64 haploid spermatids. Syncytial bridges linking spermatids within a cyst help coordinate synchronous development and elongation (25). Spermatids elongate 150- to 185-fold to reach their final length, requiring intensive reconstruction of the cytoskeleton and membrane (26, 27). Within each spermatid, microtubules arrange themselves along a pair of fused mitochondrial derivatives to form a stable zone near the nucleus, while dynamic microtubules continually extend the tail at the most distal point (27). After elongation, full-length cysts undergo individualization, in which an actin-rich individualization complex (IC) assembles around the spermatid nuclei and travels along the cyst toward the tail, condensing excess cytoplasm and unnecessary organelles into a cystic bulge that accumulates as a waste bag at the end of the cyst. As the IC migrates, it also breaks the syncytial bridges and separates spermatids into individual sperm, which then are stored in the seminal vesicles (28, 29).

Despite a detailed understanding of spermatogenesis and mechanisms of spermatid elongation, developmental processes that regulate production of sperm length diversity remain a mystery. A number of genes have been identified whose disruption interrupts elongation and is required for successful spermatogenesis (e.g., 30, 31), but fewer genetic manipulations produce sperm that differ in length but are still functional (32). During spermatogenesis, transcriptional activity is highest in late spermatogonia and early spermatocytes and lowest in late spermatids (33), confirming that post-meiotic transcription is low relative to pre-meiotic, and many gene products necessary for late stages are transcribed during earlier stages (34). It is therefore likely that genes involved in regulation of sperm length variation may be expressed at earlier stages of spermatogenesis. To identify these genes at all stages, we sequenced the transcriptomes of whole testes from males with long or short sperm derived from populations that previously underwent bidirectional selection for sperm length (20).

We found that differentially expressed (DE) genes were generally upregulated in long sperm testes, and that DE genes were enriched for Sfps and lncRNAs. To further explore the potential role of Sfps in spermatogenesis, we confirmed a putative role for two Sfps, Sex Peptide and ovulin, in sperm length variation and ruled out effects of accessory gland (AG) expression. We also found that a genetically independent population of long sperm males delays female remating relative to short sperm males, a post-mating response known to be induced by Sfp transfer during mating (35). This result suggests that Sfp expression in AG and testis is coordinated and is associated with sperm length. Our results identify a potential novel role for Sfps in regulating sperm length variation and elucidate possible mechanisms regulating natural phenotypic variation. Most Sfps are expressed both in AG and testis, they are rapidly evolving (36–39), and many are evolutionarily young (40). Moreover, the testis is a hotspot for evolution of de novo genes (41, 42). We may therefore be able to use this system to interrogate broader questions about the evolution of pleiotropy and tissue-specificity in de novo genes.

## Results

### Overview of RNAseq data

We quantified gene expression in testes from inbred isolines derived from two *D. melanogaster* populations that had been previously selected for long or short sperm (20, described in 24). After confirming sperm length differences for each isoline, we collected three replicate samples of 200 testes (from 100 males) from each isoline for a total of 12 samples. We generated RNAseq libraries from each sample, generating a total of 426.6 million mapped reads (37-51M reads/library, average of 42.6 million; **Table S1**). After filtering, we retained 10,766 annotated genes that were expressed in the testes, 9,625 of which were protein-coding. Expression profiles for all genes exhibited moderate clustering by treatment (**Fig. S1**) with a biological coefficient of variation (BCV) of 0.417, which is in line with what is expected for whole tissues (43). This BCV indicates that there was variability among samples within sperm length phenotypes but clear differences in expression profiles between phenotypes. We estimated tissue specificity using RNAseq data from 14 tissues (downloaded from FlyBase). Out of the 10,766 expressed genes, 49.7% (5347) were induced in testis and 30% (3310) had higher expression in testis compared to other tissues. The majority (3264; 61.0%) of the testis-induced genes were also induced in the accessory glands (AG), but only 9% (481) had higher expression in AG compared to other tissues. Overall, we found many genes that were highly expressed in both testis and AG, but most had the highest expression in testis. We also found a high proportion (121/176; 68.8%) of known Sfps expressed in our testes samples (33).

### DE genes between short and long sperm testes

Comparisons between short and long sperm lines revealed 317 DE genes, including 221 protein-coding genes (**Supplementary File 1**), 91 non-coding RNAs, and 5 pseudogenes. Over one third (114/317) of the DE genes were unique to *D. melanogaster*, and only 26% (82/317) were conserved across *Drosophila*. DE genes were distributed across the genome (**Fig 1, Table S2**) and the majority of DE genes (188/317; 59%) were upregulated in long sperm testes, while 129 DE genes were upregulated in short sperm testes. Across all genes, median expression levels were similar in short and long sperm testes (**Fig S2A**, Wilcoxon rank sum, FDR *p*-value = 0.58), but DE genes tended to have higher median expression in long sperm testes (**Fig S2B**, Wilcoxon rank sum, FDR *p*-value = 0.06).

**Figure 1.**
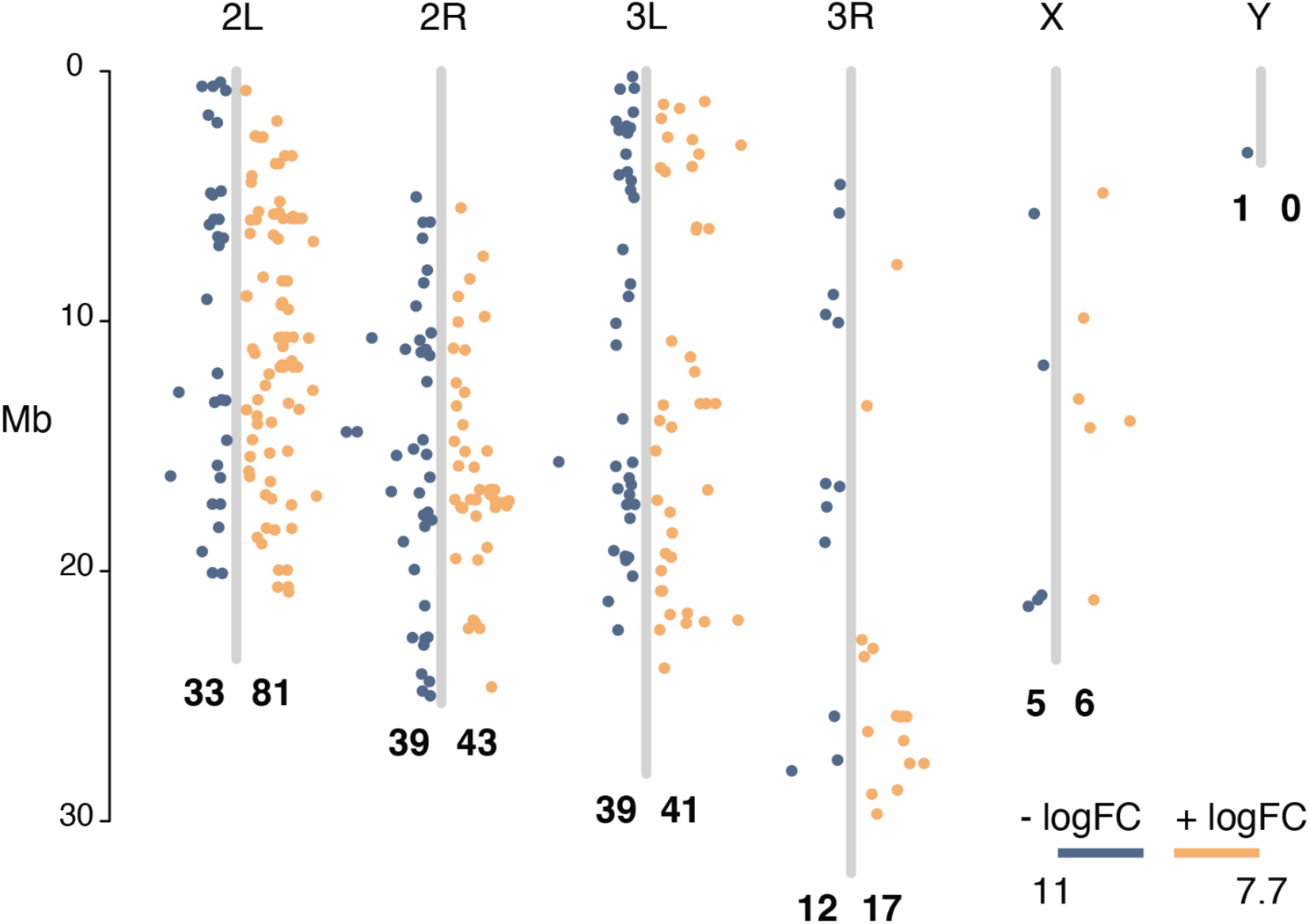
Genomic distribution of DE genes between short and long sperm producing testes. Genes with higher expression in short sperm testes (negative logFC) are blue, genes with higher expression in long sperm testes (positive logFC) are orange, and the x-axis position of each point indicates the magnitude of expression difference. Number of genes in each category per chromosome are in bold.

### DE genes are highly expressed in testes and accessory glands

DE genes had significantly higher tissue specificity relative to all genes (median ± SE τ: all genes 0.811 ± 0.002; DE genes 0.974 ± 0.006, minimum τ of DE genes 0.596). Approximately two thirds of DE genes (208/317) were induced in testis and almost one third had their highest expression in the testes relative to other tissues (65/208), most of which were genes that encoded lncRNAs (42/65, 65%; **Fig 2**, **Supplementary File 1**). Over half of the DE genes were induced in AG (162/317), and many of these had their highest expression in AG (90/162). Indeed, there were 54 known Sfps (44)differentially expressed between short and long sperm testes, all of which were induced in both the testes and the AG, but had the highest expression in the AG (**Fig 2**). There were only a handful of DE genes that were more highly expressed in other tissues (2-17 genes/tissue), and no other tissue had a high proportion of DE genes. Together, lncRNAs and Sfps comprised nearly half of the DE genes (134/317, 42%).

**Figure 2.**
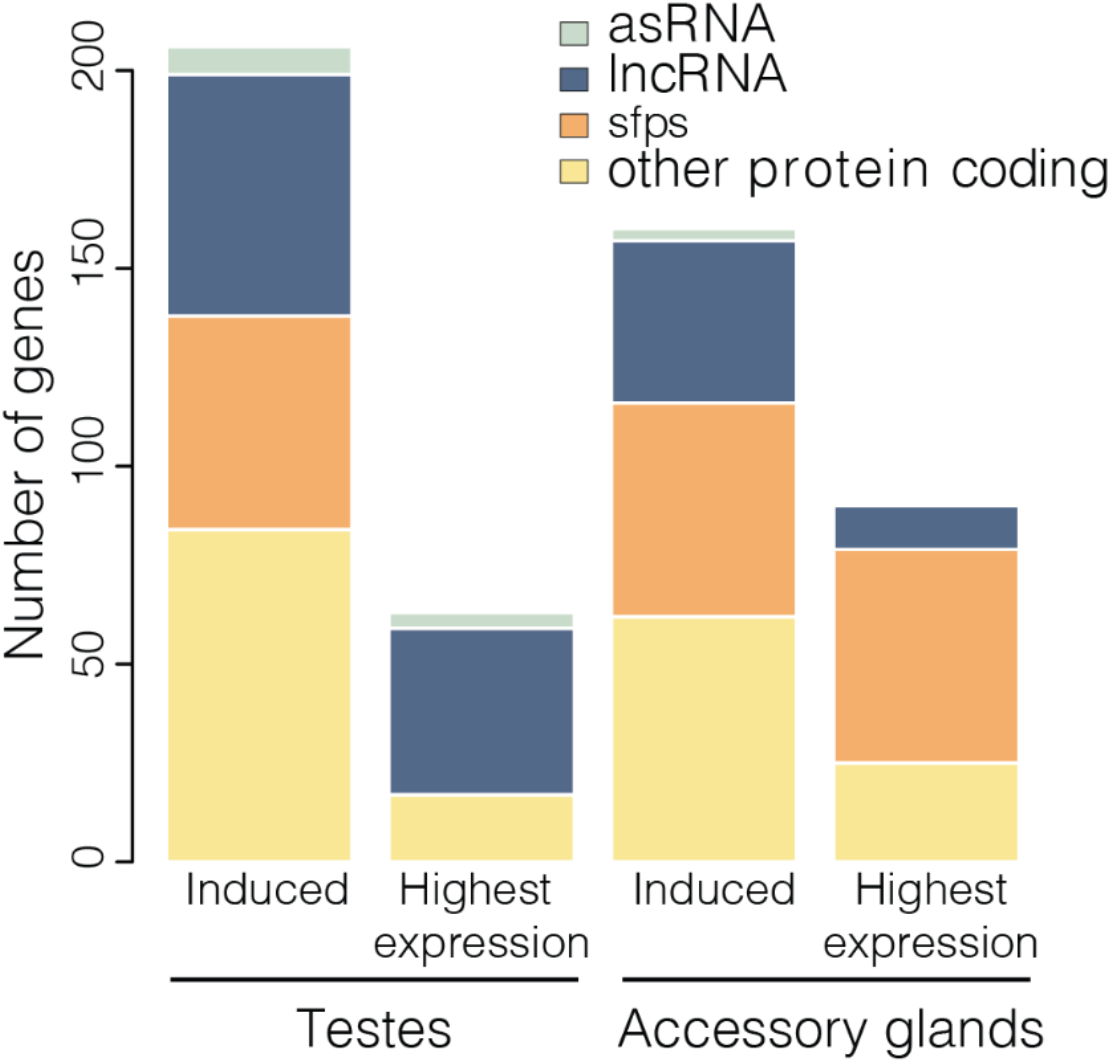
DE genes were largely testis- and AG-induced, with many Sfps and lncRNAs. Bars indicate the number of DE genes in each category, with colors indicating subcategories of genes.

### Timing of expression during spermatogenesis

To determine at what stages during spermatogenesis our DE genes are expressed, we generated a heatmap depicting stage-specific expression for 279 of our DE genes that overlapped with a previously published single-cell RNAseq dataset (33); **Fig 3**). DE genes are expressed at multiple stages of spermatogenesis and in both germline and somatic cells. Specifically, clusters of DE genes are highly expressed in epithelial cells, spermatogonia, and late spermatids. Other DE genes are also moderately expressed in cyst cells, hub cells, and spermatocytes. Of note is the small but distinct set of genes that have the highest expression in late spermatids, which is when morphogenesis and elongation occurs. Sfps and lncRNAs are differentially expressed in many cell types, with epithelial cells expressing more DE Sfps than lncRNAs, and spermatocytes and early spermatids expressing more lncRNAs than Sfps. DE genes expressed in late spermatids have a higher proportion of Sfps and lncRNAs over other gene classes.

**Figure 3.**
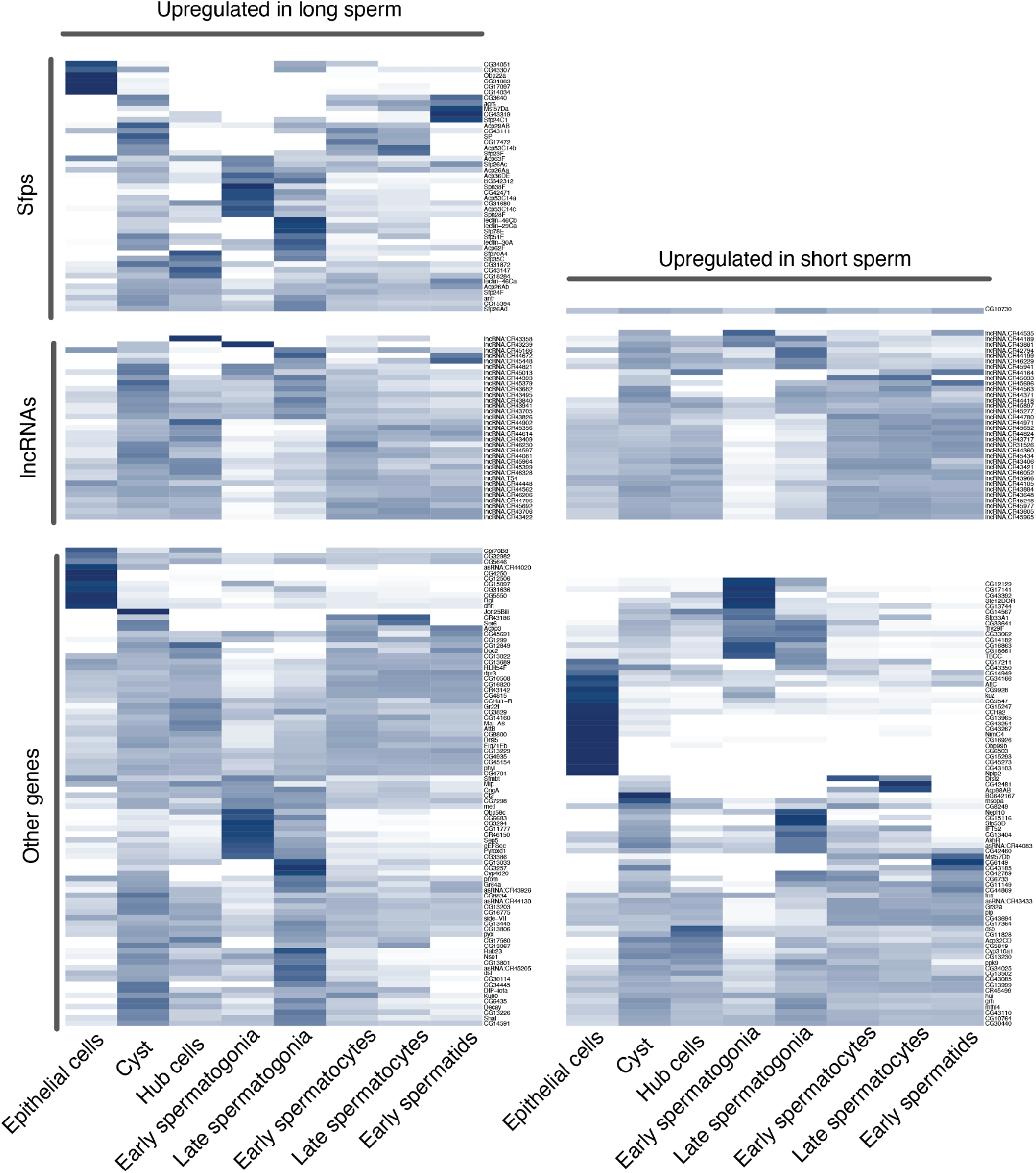
DE genes are expressed in the germline at different stages during spermatogenesis as well as in somatic cells (epithelial, cyst). Stage-specific expression derived from single-cell RNAseq dataset from (33). Only genes DE in our dataset are shown, genes are group by transcript type (Sfps, lncRNA, other) and by whether it was upregulated in long sperm (+logFC) or in short sperm (-logFC) testes.

### Gene enrichment

DE genes are most enriched for reproduction, specifically mating, sperm-related processes, and oviposition, a result likely driven primarily by the high proportion of Sfps among our DE genes (**Fig 4**). The GO terms with highest enrichment are associated with oviposition and the (often negative) regulation of female post-mating receptivity (GO:0018991; GO:0045434, GO:0046008, GO:0007621). The next highest categories are related to sperm processes including sperm competition and sperm storage.

**Figure 4.**
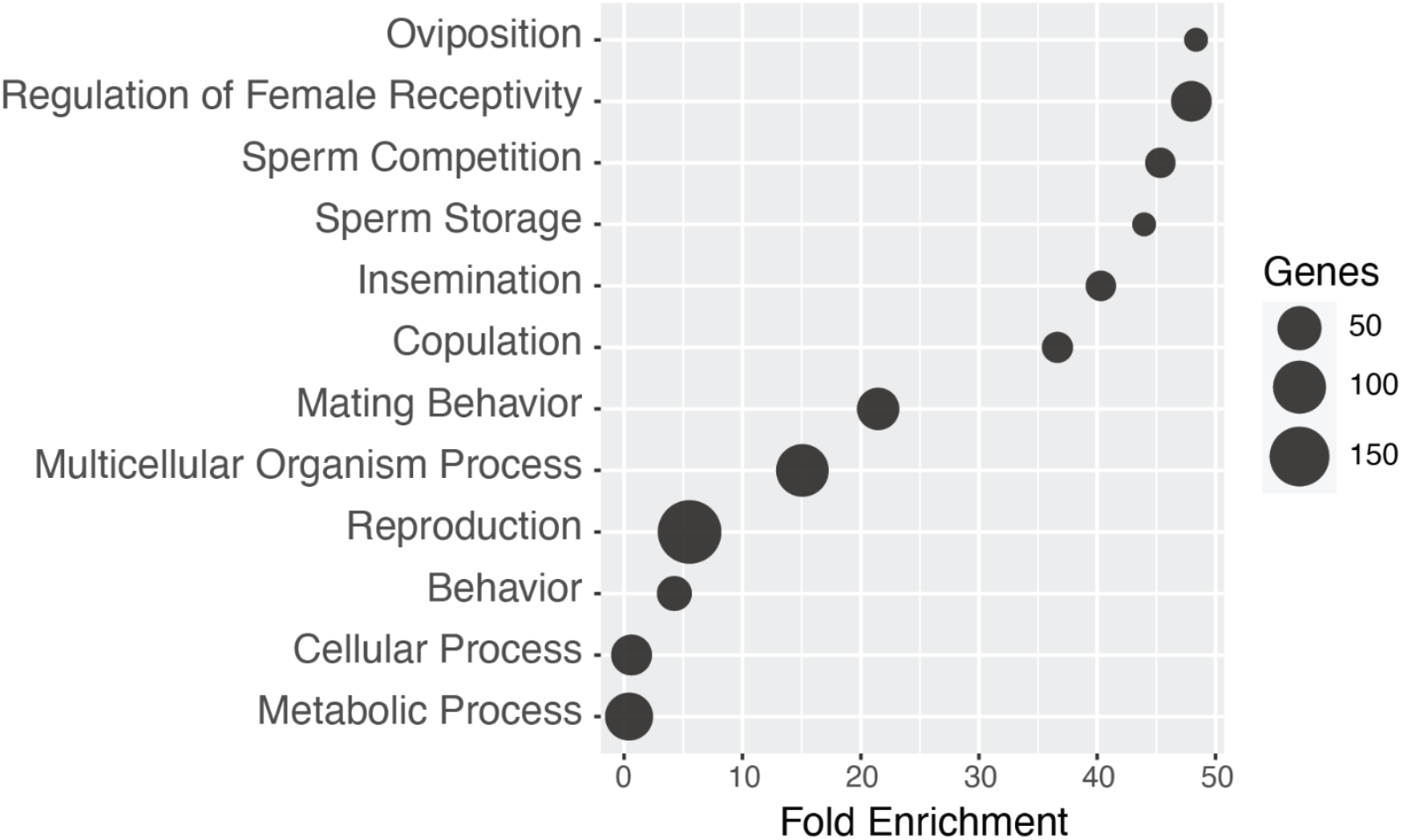
Gene ontology (GO) categories and enrichment. This bubble plot shows the top 12 GO term results from PANTHER enrichment analysis of the 317 selected DE genes. Terms are arranged in descending order by fold enrichment, and bubble size indicates the number of genes enriched for that category.

### Ruling out contamination from AG

Differential expression of Sfps in our testis samples is not likely to be due to contamination from AG tissue during dissections. Many AG-expressed genes are not expressed in our testis samples, including 55 Sfps (66) and 66 out of 74 AG-specific genes (defined as greater than low expression in AG and no/very low expression in all other tissues 45). Contamination from AG tissue would cause widespread expression of AG-expressed genes in a subset of our samples. Instead, our testis samples show expression for some AG-expressed genes, but not all, and that expression is variable depending on the gene (**Fig S3**). For example, one sample, H08C, had higher than average expression for DE Sfps, but expression of non-DE Sfps were comparable to other samples (**Fig S3**). Moreover, Sfp expression has been found in other testis expression studies. Witt et al. (2019) found Sfps that are expressed in different cell types and stages of spermatogenesis (66; **Fig 3**), and in modENCODE data, 89% of Sfps are also testis-induced (45). Finally, expression of Sfps in tissues integral to the testis, such as the epididymis and seminal vesicles, is well-documented in mammals (e.g., 46).

### Sperm size and testis length

Differential expression can result from both divergence in cellular composition or gene regulation (47). If longer sperm develop in longer testes, then expression differences between long and short sperm males could be due to overall testis size. We examined the relationship between sperm and testis length in a wild type population of *D. melanogaster* and found that they are not significantly correlated (*F*_1,43_ = 2.5673; *P* = 0.1164; **Fig S4**), suggesting that sperm length is fairly independent of testis length. This relationship was still non-significant after removing three outliers with long testes (*F*_1,40_ = 3.28; *P* = 0.078) and after applying a non-linear least squares regression to the full dataset (model: Sperm ∼ a * Testis/(b + Testis); b not significantly different from 0 with *t*_1,43_ = 1.61; *P* = 0.116). Thus, we can conclude that DE genes in our dataset are associated with sperm length and not testis size.

### Sfp knockout results in slightly shorter sperm

To interrogate the role of Sfps in sperm length variation, we performed three knockout experiments. We measured sperm in two separate genetic knockouts of Sex Peptide (SP) and ovulin (Acp26Aa), and we disrupted AG function to see if processes controlled by the AG broadly play any role in spermatogenesis within the testis. *SP* knockout males had slightly but significantly shorter sperm, by 33.81 µm in the *Δ325* x *Δ130* cross (*Χ*^2^ = 6.25, df = 1, *P* = 0.012; **Fig 5a**) but not in the reciprocal cross (*Δ130* x *Δ325; Χ*^2^ = 0.93, df = 1, *P* = 0.334; **Fig 5b**). *Acp26Aa* knockout males also had slightly but significantly shorter sperm than control males, by 72.91 μm (*Χ*^2^ = 9.89, df = 1, *P* = 0.0017; **Fig 5c**). However, any possible role of Sfps in spermatogenesis seems to be restricted to expression in testis, since knockout of AG function did not significantly alter sperm length (*F*_3,111_ = 1.38; *P* = 0.254; **Fig 5d**). Thus, while Sfp expression in testis may influence spermatogenesis, it is unlikely that AG secretions play any role in sperm length variation.

**Figure 5.**
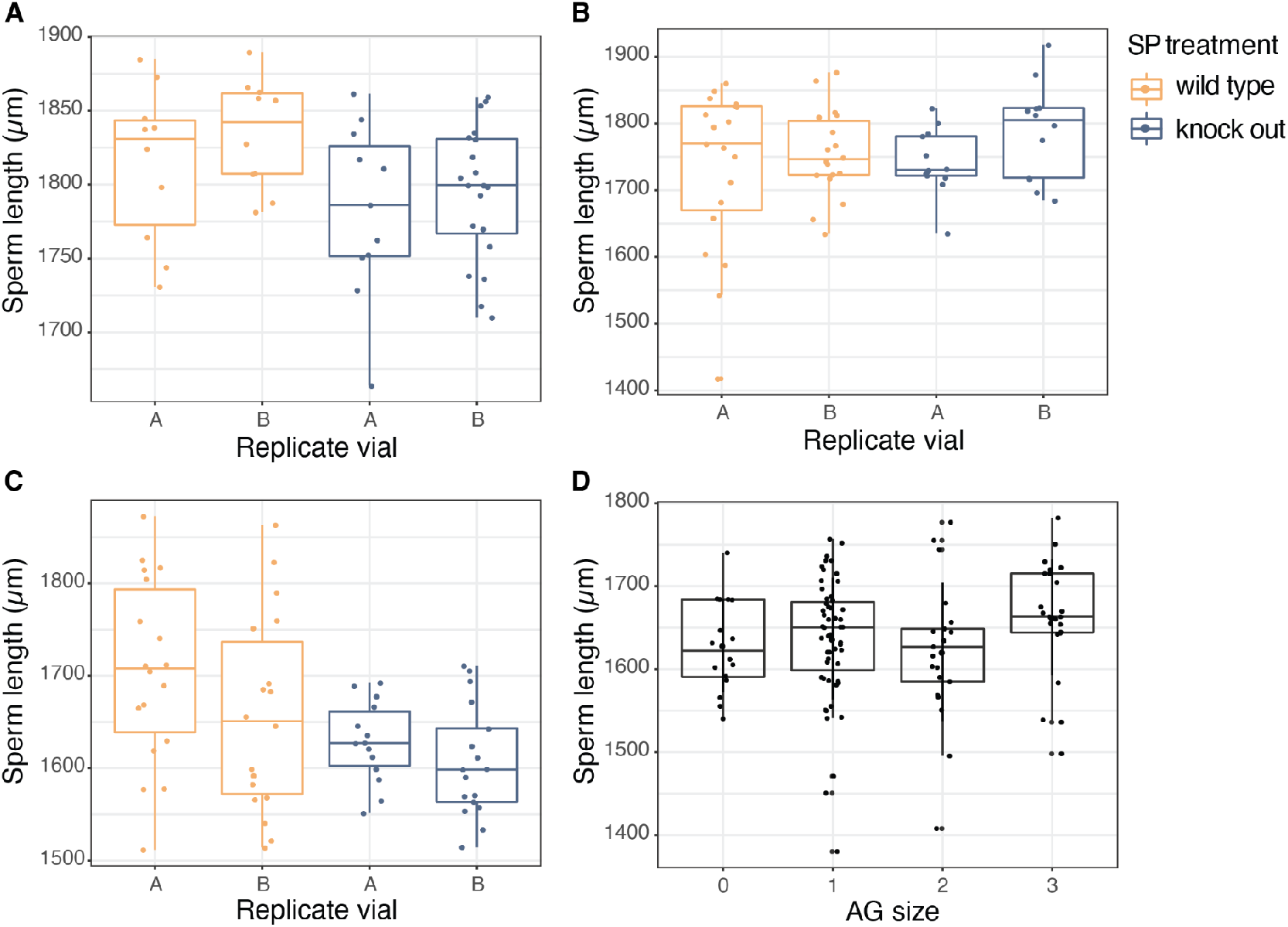
Sfp knockout in testis but not in AG may impact sperm length. Sperm are shorter in *SP* knockout males when knockout and deficiency lines are crossed in one direction (A; *P* = 0.019) but not the other (B; *P* = 0.334). *Acp26Aa* knockout males also have shorter sperm (C; *P* = 0.00075), but any role of Sfps is limited to processes in the testis, because AG knockout did not change sperm length (D; *P* = 0.169). AG size was scored from 0 (underdeveloped and non-functional) to 3 (fully developed).

### Long sperm males induce a stronger PMR

In *D. melanogaster*, Sfps are known to induce the female post-mating response (PMR), a syndrome of behavioral and physiological responses that occur after mating and include increased ovulation and oviposition, decreased receptivity to remating, and facilitation of sperm storage and release for fertilization (35). We wanted to know if increased expression of Sfps in testes producing longer sperm corresponded to higher Sfp expression in AG and thus an enhanced ability to induce the PMR. We mated wild type females first to males with long or short sperm and measured their latency to remate with a standard competitor male. Latency to remate was quantified as the time in days for half the females to remate (RT50). Long and short sperm males came from a genetically independent lineage from the RNAseq sample stocks. Males derived from a total of four recombinant inbred lines (RILs) with known sperm lengths from the Drosophila Synthetic Population Resource (48): two replicate RILs with long sperm and two RILs with short sperm. In order to assess relative fitness associated with induction of the PMR, we counted the number of progeny produced prior to remating (“prior progeny”). Because longer sperm can have an advantage in sperm competition(12, 20), we also scored paternity in progeny produced after remating, defined as P2: the proportion of progeny sired by the second (standard competitor) male.

Females mated to long sperm males took 47% longer to remate on average (0.75 days) than females mated to short sperm males. Average RT50 for mates of long sperm males was 2.35 days, compared to 1.4 days for mates of short sperm males (RIL 22059: 2.2 days, 22096: 2.5 days, 22097: 1.8 days, 22125: 1.4 days; **Fig 6a**). However, this delay in female remating was not enough to result in more prior progeny (*Χ*^2^ = 0.404, df = 1, *P* = 0.525). Long sperm also did not provide an advantage in sperm competition, because there was no difference in paternity success between long sperm and short sperm males (*z* = −0.218, *P* = 0.828).

**Figure 6.**
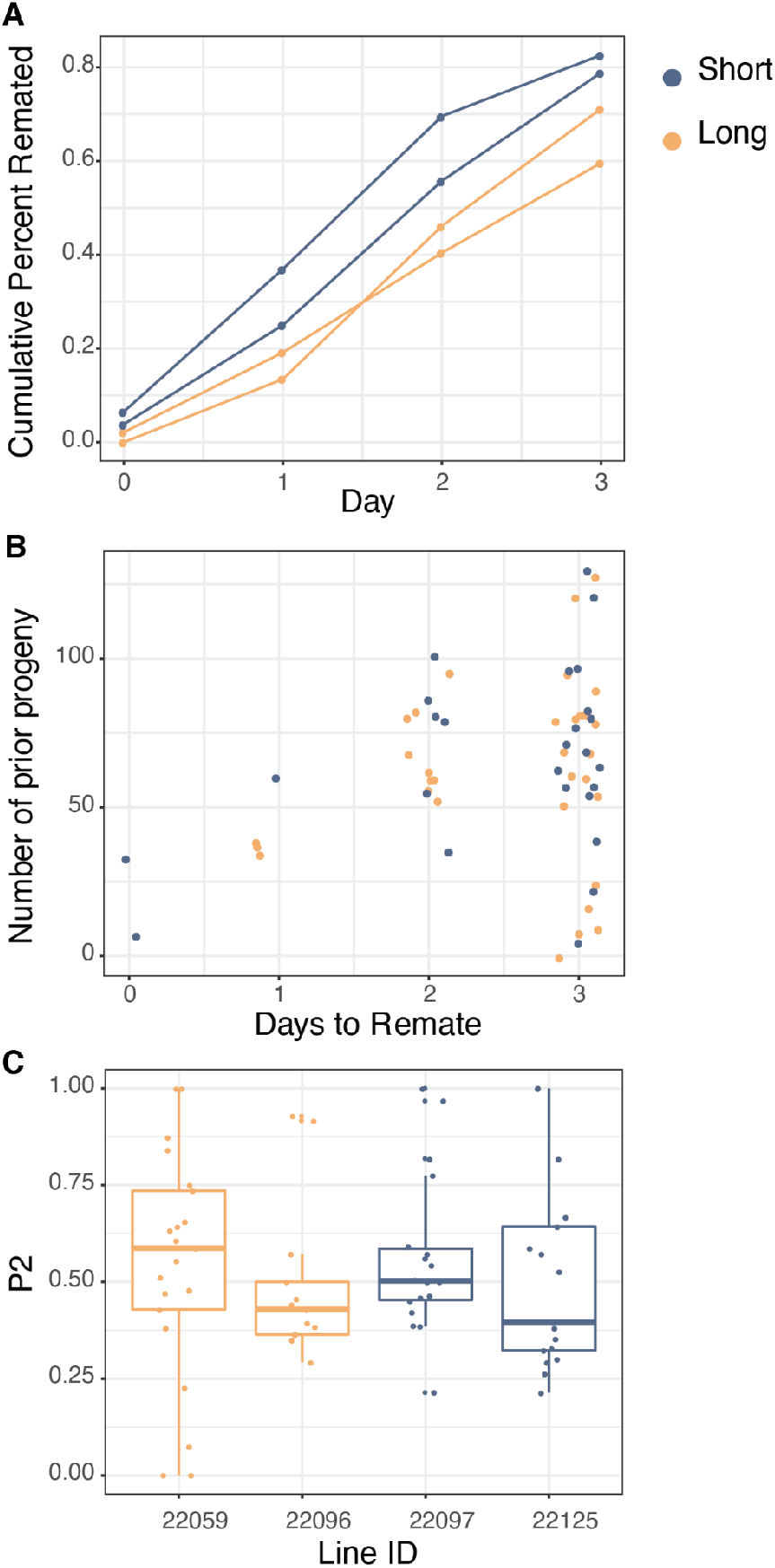
Long sperm males delayed female remating but did not have an associated increase in fitness. (A) Females mated to long sperm males delayed remating relative to mates of short sperm males, but (B) this did not translate into more prior progeny (*P* = 0.525). (C) Long sperm males also did not have higher paternity success against a standard competitor male (*P* = 0.828). Long sperm RILs are in yellow, short sperm in blue.

## Discussion

In this study, we set out to find candidate genes that may control natural variation in sperm length in healthy males exhibiting a typical range of phenotypic variation. Our DE genes were enriched for lncRNAs and Sfps, and sperm length in SP and ovulin knockouts was slightly but significantly shorter than wild type controls. We ruled out the possibility that the Sfp effect was due to cross-communication from AG-expressed Sfps by showing that sperm length was not affected by knocking out AG function. Next, we asked if higher expression of Sfps in long sperm testis was also associated with higher expression of Sfps in AG. We assayed males with known sperm lengths from a genetically independent synthetic population for their ability to induce the female post-mating response. We found that long sperm males delay female remating for longer than short sperm males, but this delay did not yield significant advantages in the number of prior progeny or paternity success. Thus, we have found that male reproductive success is mediated by coordination of two distinct tissues in the male reproductive tract that each contribute different and essential components of the ejaculate. However, despite abundant evidence for the importance of both Sfps and sperm length in post-copulatory sexual selection, the fitness advantages of such coordination within a competitive context are not clear. It is possible that although the fitness advantage is undetectable in our dataset under lab conditions, it could be enough to significantly impact the evolutionary trajectory of a large wild population. Selection on the amount of Sfps transferred may be weak if a threshold amount is sufficient to induce the female PMR. As a result, natural variation in Sfp transfer appears to cause differences in female remating latency but not at a level that will affect fecundity and paternity success.

In *D. melanogaster*, 176 Sfps have been confirmed that are produced in the male reproductive tract and transferred to females (44). They are a diverse set of proteins, including small signaling peptides, proteases, protease inhibitors, lectins, anti-oxidants, odorant-binding proteins, and large pro-hormones (35, 44, 49). Some of these molecules enter the female reproductive tract in a physiologically inert state and become activated upon proteolytic cleavage (50–53). After mating, Sfps induce changes in female physiology, behavior, gene expression, and morphology, including increased oviposition, sperm storage, feeding, and reduced receptivity to remating (35, 49, reviewed in 54–57). These changes can occur over the course of hours or days and are collectively referred to as the post-mating response (PMR). Sfps are also subject to sexually antagonistic coevolution, such that females that mate more die faster (58), especially when not allowed to coevolve with males under polyandry (59). Some Sfps have been shown to contribute to competitive fertilization success (60, 61), and like other reproductive proteins, tend to evolve rapidly (37, 40, 62–64). Although historically attributed to positive selection, recent analysis that incorporates intraspecific polymorphisms has found that this rapid evolution may not always be adaptive in nature but rather a consequence of relaxed selection associated with the evolution of sex-biased expression (65, 66). In any case, rapid evolution of Sfps means that they are often not conserved across more distantly related species. It remains to be seen whether upregulation of Sfps is a consistent hallmark of longer sperm in other species and whether it is dependent on species-specific sperm lengths. It is possible that developmental regulation of intraspecific variation in sperm length depends on whether sperm length is 2 mm or 2 cm.

We selected SP and ovulin for further investigation, because they are among the best-studied Sfps in *Drosophila*. SP is a small 36-aa peptide that binds to sperm tails in the SR and is slowly released over several days, mediating the decline of female receptivity that characterizes the long-term PMR (67). In testis, SP expression is highest in spermatocytes and cyst cells (33), so they may be important for downstream processes, but SP localization in testis is unknown. One question that arises from our results is whether SP binds to sperm during spermatogenesis in addition to binding after ejaculation in the female reproductive tract (67, 68). It is also possible that longer sperm can bind more SP, which could affect the PMR, but both of these factors remain to be demonstrated. More recently, SP was shown to be required for assembly and disassembly of lipid-rich microcarriers that mediate transfer of seminal fluid components in the ejaculate (68). Exosomes in mouse testis are transporters of non-coding RNAs that are required for sperm maturation (69) and normal embryonic development (70), but the function of testis-based exosomes in *Drosophila* spermatogenesis is unknown. Ovulin is a large 264 aa pro-hormone (71) that is cleaved in the female reproductive tract into two products that, along with SP and other Sfps, stimulate oviposition as part of the short-term PMR (72). It is possible that ovulin undergoes testis-specific cleavage to serve a different function in that tissue. It is upregulated by over five-fold in late-stage spermatids (33), suggesting it is important during spermatid elongation. Future research should focus on localizing SP, ovulin, and other Sfps in the testis as well as conducting additional genetic manipulation experiments to further elucidate the roles Sfps play in spermatogenesis.

Among our DE genes, over half are induced in AG, including nearly one third of known Sfps. These genes are typically expressed primarily or exclusively in the male reproductive tract with the highest and next-highest expression levels in the AG and testis, respectively. We know of no other examples of such closely coordinated gene expression between two tissues, and it is unclear why this coordination has evolved or how it is accomplished mechanistically. Though both tissues are part of the male reproductive tract, testis and AG are morphologically, physiologically, and developmentally distinct. AG is epithelial in origin and arises during metamorphosis from the male genital primordium of the genital disc (73), while testes (and ovaries) develop during embryogenesis from mesodermal cells that differentiate into somatic gonadal precursors and then gonads (74). In the testis, gene expression is regulated at different spermatogenic stages by shifting suites of transcriptional regulators including testis-specific meiotic arrest complex (tMAC) and testis-specific TBP-associated factors (tTAF), which prevent meiosis until terminal differentiation genes have sufficiently accumulated in spermatocytes (75). In AG, *HR39* (76), *dve* (77), and *prd* (78) are each required for Sfp expression and male fertility. They are all also expressed in the testis, and *dve* is upregulated 5-fold in late spermatid cysts (33). *HR39* knockout also decreases expression of genes in the testis (76), pointing to one potential mechanism for testis-AG coordination. It is not out of the question that there can be gene network interactions among different tissues of the reproductive tract (e.g., 79), but the fact that AG ablation doesn’t affect sperm length suggests that in this case, coordination is accomplished by other means. Characterization of regulatory mechanisms for Sfps compared to truly AG- or testis-specific genes may yield additional insights.

LncRNAs comprised 25% of our DE genes, and though less well-annotated than the Sfps, they may be just as important in regulating sperm length. LncRNAs generally regulate gene expression in many different tissues via diverse mechanisms (80). Many lncRNAs are testis-specific, expressed in all stages and cell types (81), and are differentially expressed in association with male fertility (82). In *Drosophila* testis, most lncRNAs with stage-biased expression are upregulated post-meiotically during elongation, suggesting they play a significant role in sperm morphogenesis and maturation (83). Indeed, functional characterization of a testis-specific lncRNA in *Drosophila* resulted in significant defects during late spermatogenesis (84). Another knockout screen found fertility defects in 31% of testis-specific lncRNAs examined (85), including three DE genes in our dataset (*CR43633*, *CR44344*, and *CR44371*). Next steps should characterize mechanisms of the male subfertility phenotype for these genes to better understand their roles in spermatid elongation and maturation.

Reproduction is already a complicated affair, and knowing that sperm length is associated with Sfp expression adds another layer of complexity. Competitive reproductive success is a function of male traits, female traits, and their interactions over the course of the reproductive process from mating to fertilization. During mating, sperm and seminal fluids mingle in the female’s bursa (68) and enter the SR, where sperm physically displace resident sperm from previous matings back into the bursa (86, 87). Displacement continues until the female ejects excess sperm from the bursa, and the timing of this ejection influences the proportion of second-male sperm remaining in the SR for fertilizations (87, 88). Several hours after ejection, females begin to ovulate, sperm are released from the SR according to a fair raffle (86), and eggs are fertilized in the bursa. Sfps are known to influence several aspects of this process, and displacement is also a function of SR length and the difference in sperm lengths between the two males (12). Sperm length and SR length are positively genetically correlated, as are SR length and remating rate (22). Here, we showed that long sperm are associated with a delay in female remating, but these two traits are not genetically correlated (22). Delayed female ejection is also associated with long sperm (12), suggesting that Sfps may influence the timing of ejection as well, which would further amplify competitive fertilization success for long sperm males.

All these coordinated advantages of sperm length, Sfps, delayed remating, and delayed ejection suggest that long sperm males should have an ultra advantage in sperm competition, but this does not seem to be the case. In this study, long sperm males did not sire more progeny either before remating (prior progeny) or after remating (paternity success). Male competitive fertilization success is non-transitive, such that one male is never successful against all other males when mating with all other females. Rather, male (and female) success is a function of many interacting factors, including sperm length, SR length, sperm numbers, female size, and Sfps (12, 89). In other words, the fitness value conferred by a phenotype depends largely on other interacting phenotypes, and these direct and indirect genetic effects have implications for the direction and rate of phenotypic evolution (90). The end result is that selection on sperm length in *D. melanogaster* is not simply directional (or stabilizing). This context-dependent selection complicates the fitness landscape and makes it less predictable, while also likely maintaining high rates of sperm length variation within populations.

The genetic covariance of sperm and seminal fluid components of the ejaculate is not necessarily surprising. After all, quantitative genetic theory predicts that functional and developmental integration of traits will lead to their genetic integration, which in turn leads to evolutionary integration (91). Sperm and Sfps are arguably not developmentally integrated, based on our inability to generate a sperm length phenotype after knockdown of AG function. However, sperm and Sfps are certainly functionally integrated, given that Sfps are required for normal sperm function (92). Recently, genetic covariance has also been found in *D. bipectinata* between male sex combs (which help grasp the female during mating) and Sfp expression, leading to enhanced competitive fertilization success for males with larger sex combs (93). This coordination of Sfps with a non-genitalic copulatory trait suggests that evolutionary integration could be expected for an even wider range of sexually selected traits. Indeed, such evolutionary integration between a male ornament and female preference is required for Fisherian runaway selection (but see 94, 95) and has been found for trait-preference/perception systems in *Drosophila* (22, 96, 97), cricket (98), dung beetle (99), and medaka (100). The molecular mechanisms and evolutionary consequences for integration of sexually selected traits are not well-understood (101), but the *Drosophila* ejaculate and female reproductive tract can be used as a well-characterized model system in which to further explore these questions. Moreover, because the testis is a hotspot for de novo gene evolution (41), this system can also be harnessed to test hypotheses about how pleiotropy evolves over the course of a new gene’s developmental trajectory. One question asks whether newer genes are less likely to be functionally integrated with AG and whether older sex-biased genes have evolved specificity of expression in non-testis tissues. Answers to these questions will inform our understanding of the evolution of novelty and complexity in the context of sexual selection.

## Materials and Methods

### RNAseq libraries and analysis

We used inbred isolines derived from two *D. melanogaster* populations that had been previously selected for long or short sperm (20, described in 24). Briefly, the original populations underwent 17 generations of selection for sperm length, followed by approximately 300 generations of random mating. They were then inbred through 10 generations of full-sibling mating, resulting in a panel of isolines with short or long sperm. To confirm the differences in sperm length for each isoline, approximately five sperm from at least four males (range 4–8 sperm, average: 5.56 sperm) were measured (see 24). We selected two isolines with long sperm (H08, H20) and two with short sperm (L08, L17) and maintained breeding vials at 23°C with a 12:12 light:dark cycle on sugar-yeast-agar diet in vials with approximately 1.5 cm^3^ medium supplemented with live yeast.

We collected three replicate samples of 200 testes (from 100 males) from each isoline for a total of 12 samples. We collected males within 24 hours of eclosion and aged them 4 to 6 days in food vials with live yeast, at densities of up to 20 males per vial. We dissected testes under ether anesthesia with fine Dumont tweezers (Ted Pella cat. no. 505) into a droplet of sterile Grace’s physiological insect medium. We washed testes in fresh medium, transferred them to 200 μl of Trizol and froze them at −80°C until RNA extraction. We isolated total RNA using a low sample volume Trizol-chloroform extraction (protocol from 102) and quantified RNA using an Agilent Bioanalyzer 2000. We omitted two low quality samples and all but one of the remaining 10 samples had RIN > 7.0. Total RNA was sent to the Huntsman Cancer Institute at the University of Utah, which prepared Illumina TruSeq Stranded mRNA libraries with PolyA selection and rRNA depletion. Libraries were pooled and sequenced on an Illumina HiSeq 2000 (PE, 100bp).

We trimmed adaptors and removed low quality reads using TRIMMOMATIC v0.39 (103). We mapped reads to the *D. melanogaster* genome (BDGP6.28) using HISAT2 v2.2.0 (104) with default settings. We counted the number of reads that uniquely mapped to annotated genes (Ensembl release 100) using FEATURECOUNTS v1.4.4 (105). We analyzed gene expression using BIOCONDUCTOR v3.0 package edgeR v3.30.3 (106) in R v4.0.1 We normalized our data using the scaling factor method and restricted our analysis to genes with a minimum expression of FPKM > 1 in at least four samples. For all analyses, we tested alternative normalization methods (weighted trimmed mean of M-values) and found qualitatively similar results. We fit our data with a negative binomial generalized linear model with Cox-Reid tagwise dispersion estimates (107). To evaluate differential expression, we used likelihood ratio tests, dropping one coefficient from the design matrix and comparing that to the full model. For all of our results we used a p-value adjusted for a false discovery rate (FDR) of 5% (108).

We quantified tissue specificity using RNAseq tissue expression data from FlyBase (gene_rpkm_report_fb_2020_04.tsv) for fourteen tissues (see **Supplementary File 1**). We defined a gene as testis-induced if its expression was greater than twice its median expression in other tissues. We estimated tissue specificity (τ) following the recommendations of Liao and Zhang (109). The τ value ranges from 0 to 1 with higher values indicative of expression restricted to one or a few tissues (109–111). We used the PANTHER Gene Ontology (GO) resource (112) to perform an over-enrichment test on all 317 differentially expressed (DE) genes between long and short sperm. Specifically, we performed a Fisher’s exact test with FDR correction, comparing our gene list with a *D. melanogaster* reference set from the PANTHER database (113, 114).

### Testis size and sperm length

We tested the relationship between sperm and testis length in a wild type population of *D. melanogaster* (LHm; 115). This stock was reared on sugar-yeast-agar medium sprinkled with live yeast at room temperature (∼23°C) with ambient light. We collected 45 newly eclosed virgin males and aged them for 3-5 days in same-sex vials at densities of up to 20 per vial. We anesthetized males with ether and isolated sperm from one testis and mounted the other testis for measurement. To obtain sperm, we dissected seminal vesicles into a large droplet of 1X phosphate-buffered saline (PBS) on a glass slide and ruptured the tissue to release motile sperm. We dried the droplet down at 50-60 °C and fixed the sperm in 3:1 methanol:acetic acid, mounted in glycerol, and sealed the coverslip with nail polish. We visualized sperm on a Nikon Ni-U upright light microscope at 100X or 200X magnification under darkfield, captured images with an Andor Zyla 4.2 camera, and measured sperm length using the segmented line tool in ImageJ (https://imagej.nih.gov/ij/), adjusting for scale at different magnifications. We measured 1-7 sperm per male, with an average of 4. These sample sizes are standard (e.g., 24, 32) and sufficient to capture variation among males (**Fig S5**). To measure testis size, we dissected a testis with attached seminal vesicle using fine forceps in 1X PBS and transferred the tissue to 40 μl of PBS, mounted under a cover slip, imaged immediately at 100X under phase contrast, and measured using the segmented line tool in ImageJ. We assessed the relationship between testis length and sperm length using both linear regression (lm) and nonlinear least squares regression (nls) in R v3.4.3.

### Sfps and sperm length

All stocks and crosses were maintained for at least two prior generations on sugar-yeast-agar medium sprinkled with live yeast at 23 °C with 12:12 light:dark cycle. We generated *SP* null mutant males by crossing the *SP* knockout line *Δ325/TM3, Sb, ry* with an *SP* deficiency line, *Δ130/TM3, Sb, ry* (116), in both directions (*Δ325* female x *Δ130* male; *Δ130* female x *Δ325* male). Experimental knockout males were identified by wild type *Sb+* phenotype, while control siblings were *Sb*. To knock out ovulin, we crossed the mutant stock *Acp26Aa1* (117) with a chromosomal deficiency mutant missing a 140 kb region on chromosome 2L that includes *Acp26Aa* (“Df(2L)Exel6014”; Bloomington Drosophila Stock Center #7500;, 118). This knockout cross was also set up in both directions, but only *Acp26Aa* (female) x *Df(2L)Exel6014* (male) yielded enough progeny. Experimental knockout *Cy+* males were compared with control *Cy* siblings. Both the SP and ovulin knockout crossing schemes allow us to examine the knockout phenotype while minimizing associated genetic effects that may have accumulated within the individual lines.

The AG knockout was achieved by inducing strong endoplasmic reticulum stress within the AG, inhibiting maturation and full AG function (119). This was done by driving UAS-mediated expression of the misfolded protein associated with allele *Rh1G69D* (120) with the AG-specific *prd-GAL4* driver (78). The ensuing unfolded protein response (UPR) resulted in AGs that were small, underdeveloped, and empty. Progeny included knockout siblings (*Cy*+, *Sb*+) and control siblings expressing either TM3 balancer (*Sb*; *prd-GAL4*; Bloomington Drosophila Stock Center #1947) or CyO (*Cy*, *UAS-Rh1G69D*). Because not all knockout siblings had nonfunctional AGs, we examined the relationship of sperm length with AG phenotype directly, rather than with *Sb*+ or *Cy*+ phenotypes. AG phenotype was scored on a scale from 0 (underdeveloped and non-functional) to 3 (fully developed and wild type).

For all knockouts, adult males were collected from two replicate vials (A and B) within 24 hours of eclosion, aged 5-7 days, and sperm were collected, prepared, and measured as described above. Numbers of sperm measured per male varied from 1 to 13, with an average of 6.8 to 7.7 sperm per male. All stocks were generously provided by Dr. Mariana Wolfner, including *Rh1G69D* with permission from Dr. Hyung Dong Roo.

All statistical analyses were performed in R v3.6.3. Depending on the dataset, outliers below 1000 or 1300 μm and above 2000 or 2300 μm were presumed to be broken sperm or human error and removed, resulting in omission of 1 to 34 measurements per dataset. SP and ovulin knockout data were analyzed using linear mixed-model regression (lmer in the package lme4) fitted by restricted maximum likelihood (REML). The model consisted of treatment (knockout or control) as a fixed effect, and replicate vial and male as random effects nested within treatment (model: sperm length ∼ treatment + (1 | male : replicate : treatment)). Significance was estimated using a Type II Wald chi square test implemented with Anova in the car package. AG knockout data were analyzed using simple ANOVA of sperm length across AG sizes using aov.

### Sperm length and the post-mating response

Wild type females were from an M3 wild type stock collected from Silver Spring, Maryland by one of the authors (MKM) in 2018. Long and short sperm males came from a total of four recombinant inbred lines (RILs) with known sperm lengths from the Drosophila Synthetic Population Resource (48): two replicate RILs with long sperm (RIL ID no. 22059, 22096) and two RILs with short sperm (ID no. 22097, 22125) that were originally phenotyped as part of another study in 2019. The standard competitor male derived from a Canton-S stock with a protamine-GFP construct that is expressed in sperm heads as well as an external GFP eye marker for scoring paternity (32). All stocks new to the lab were reared for at least two prior generations on yeast-sugar-agar medium at 23 °C sprinkled with live yeast at moderate densities.

We remeasured sperm lengths in the RILs to verify their phenotype, by dissecting and preparing sperm samples as described above, measuring 5 sperm per male for 3 males per RIL. Sperm length distributions for long and short lines remained non-overlapping, with long lines averaging 1933 ± 19.7 μm and 1874 ± 13.0 μm, and short lines averaging 1555 ± 90.4 μm and 1555 ± 15.2 μm. Average sperm length for the standard competitor Canton-S GFP stock was 1854 ± 23.4 μm, and average SR length of the M3 stock was 2515 ± 47.2 μm (**Fig S6**).

To assess remating latency, M3 virgin females (aged 3-4 days post-eclosion) were aspirated (without anesthesia) into individual food vials supplemented with live yeast and left overnight to acclimate. The next morning, a single male from one of the four RILs (aged 3 days) was aspirated into each female vial. If copulation occurred, the male was removed and discarded, and a standard competitor male (aged 2-6 days) was introduced into the vial (Vial 1). Females were provided with a daily four-hour opportunity to mate and remate over four consecutive days. We noted the time of male introduction, as well as start and end times of first and second copulations. When females remated, the second male was discarded, and females were transferred to a fresh food vial (Vial 2), where they laid eggs for seven days. Progeny were scored for paternity from Vial 2, and prior progeny were counted from Vial 1. The experiment ended when at least 50% of females mated to each RIL remated. Remating rate for each RIL was quantified as the time in days until 50% of females remated (“RT50”). ImageJ was used to precisely measure RT50 for each RIL from its cumulative remating curve.

We used mixed model regression to test for an effect of sperm length on the number of prior progeny using linear mixed-model regression (lmer in the package lme4) fitted by restricted maximum likelihood (REML). The model consisted of sperm length (long or short) as a fixed effect and two random effects: replicate RIL nested within treatment and number of days to remate (model: prior progeny ∼ sperm length + (1 | days to remate) + (1 | sperm length : RIL)). Significance was estimated using a Type II Wald chi square test implemented with Anova in the car package.

Paternity scoring with the Canton-S GFP stock was complicated by partial loss of the transgene, along with weak or lost GFP eye signal. To work around this challenge, we limited paternity scoring to sons and examined GFP-protamine expression in testes. If at least one son expressed GFP-protamine, we scored paternity in all sons from that female. This approach allowed us to score paternity for 13 to 21 families in each RIL treatment. To test for differences in P2 (proportion of progeny sired by the second male), we used logistic regression with a logit link function and binomial error distribution (after ensuring no overdispersion in the data), implemented using glm in R v3.6.3.

Stocks used in all experiments are listed in **Table S3**.

### Data Availability

The data reported in this paper are available through the National Center for Biotechnology Information Sequence Read Archive under accession number XXXXXXX and on Dryad under access number XXXXXX.

## Acknowledgments

We are grateful to MF Wolfner for stocks and advice, S MacDonald for DSPR stocks, S Wigby and B Hopkins for helpful conversations, and many undergraduate researchers for assistance with data collection, especially I Abdullah, E Bansal, V Barone, L Cornell, I Diwani, O Fanshawe, T Huynh, S Kujawski, K Markollari, L Morse, O Saadi, and A Stathopoulos. This work was supported by a National Science Foundation (NSF) grant DEB-1257859 to MKM, a University Faculty Fellowship from George Washington University and ELL was supported by NSF DEB-1557059.

## Supplemental Files

**Supplementary File 1. Summary of DE genes between short and long sperm producing testis.** This summary file (.csv) contains each gene’s FlyBase ID, Gene name, Chromosome, Gene type (i.e., protein coding, ncRNA etc), logFC between short and long sperm producing testes, and FDR corrected p-value. It also contains a summary of expression in typical *D. melanogaster* using data from FlyBase, including the tissue specificity index (tsi), whether the gene was expressed in typical *D. melanogaster* testes, expressed in short sperm, long sperm, induced or had the highest expression in typical *D. melanogaster* testes, male accessory glands, what type of ncRNA and whether the gene is a characterized Sfp.

**Figure S1.**
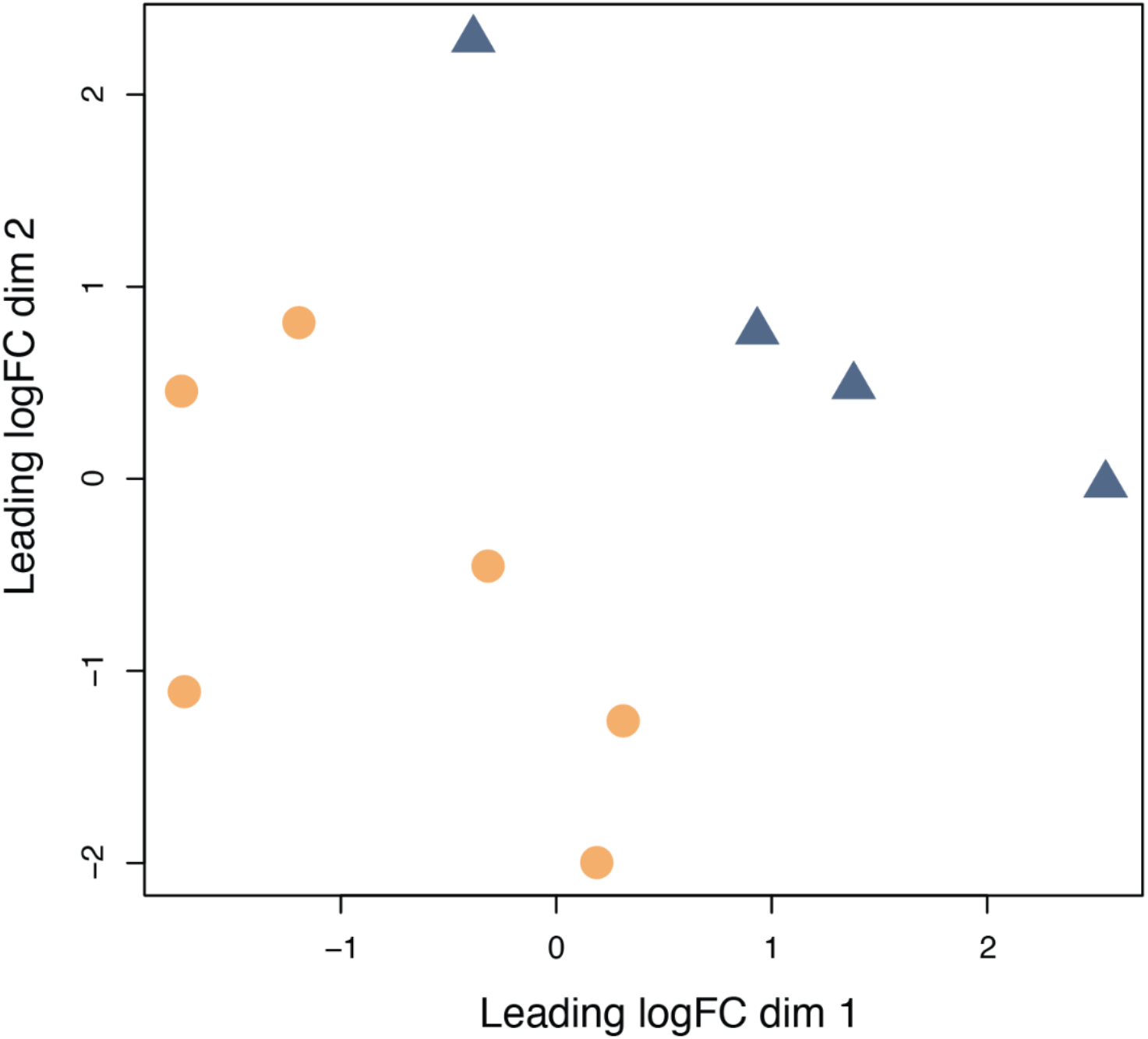
Clustering of gene expression profiles. Multidimensional scaling plots (MDS) of the Euclidean distance among gene expression profiles. Distance approximates the typical log2 fold changes between samples for the 500 genes with the greatest expression differences among treatments. Samples from short sperm testes are blue triangles and samples from long sperm testes are orange circles. There was moderate variation among samples (biological coefficient of variation = 0.417), which is overall consistent with other whole-tissue gene expression profiles (47).

**Figure S2.**
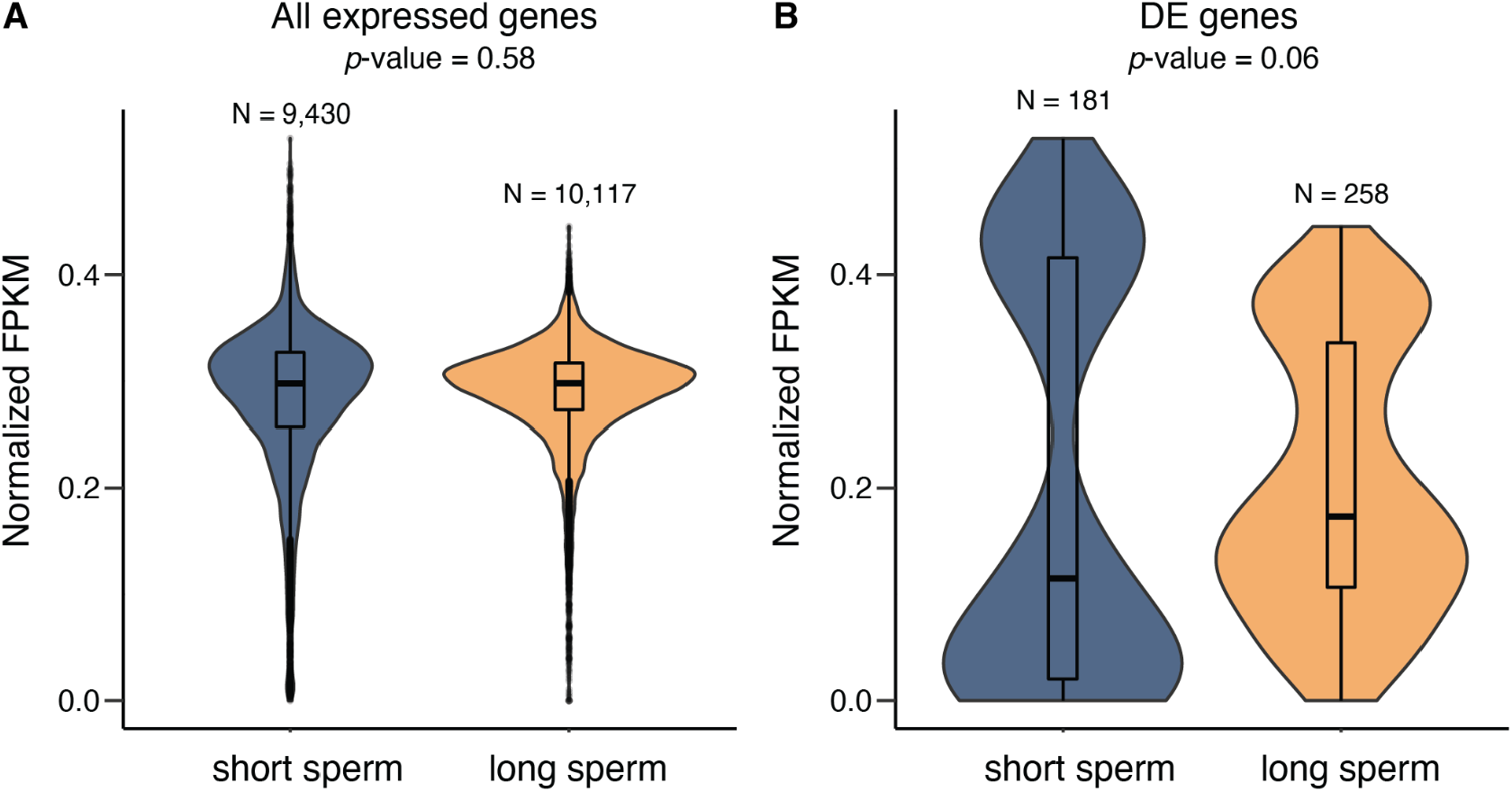
Normalized expression for A) all expressed genes and B) DE genes in short and long sperm producing testes. These violin plots show the probability of density of expression values with wider portions indicating a greater number of genes with that expression value. The boxplots at the center of each violin plot are the median expression and quartiles. We tested for differences in median expression between short and long sperm producing testes using a Wilcoxon Rank Sum test with FDR corrected p-values.

**Fig. S3.**
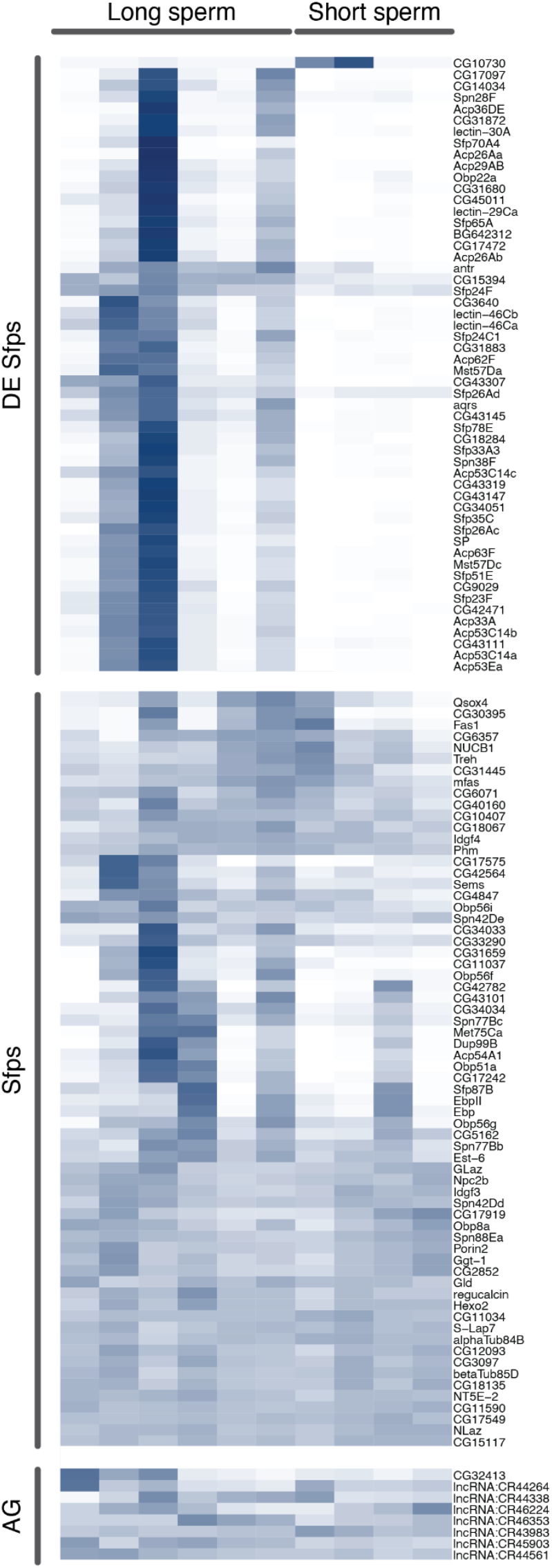
AG-expressed genes are expressed to varying degrees within and across samples, suggesting DE of Sfps is not due to contamination. Heat map showing gene expression across long (N = 6) and short (N = 4) sperm samples for Sfps that are DE and non-DE as well as genes thought to be AG-specific.

**Figure S4.**
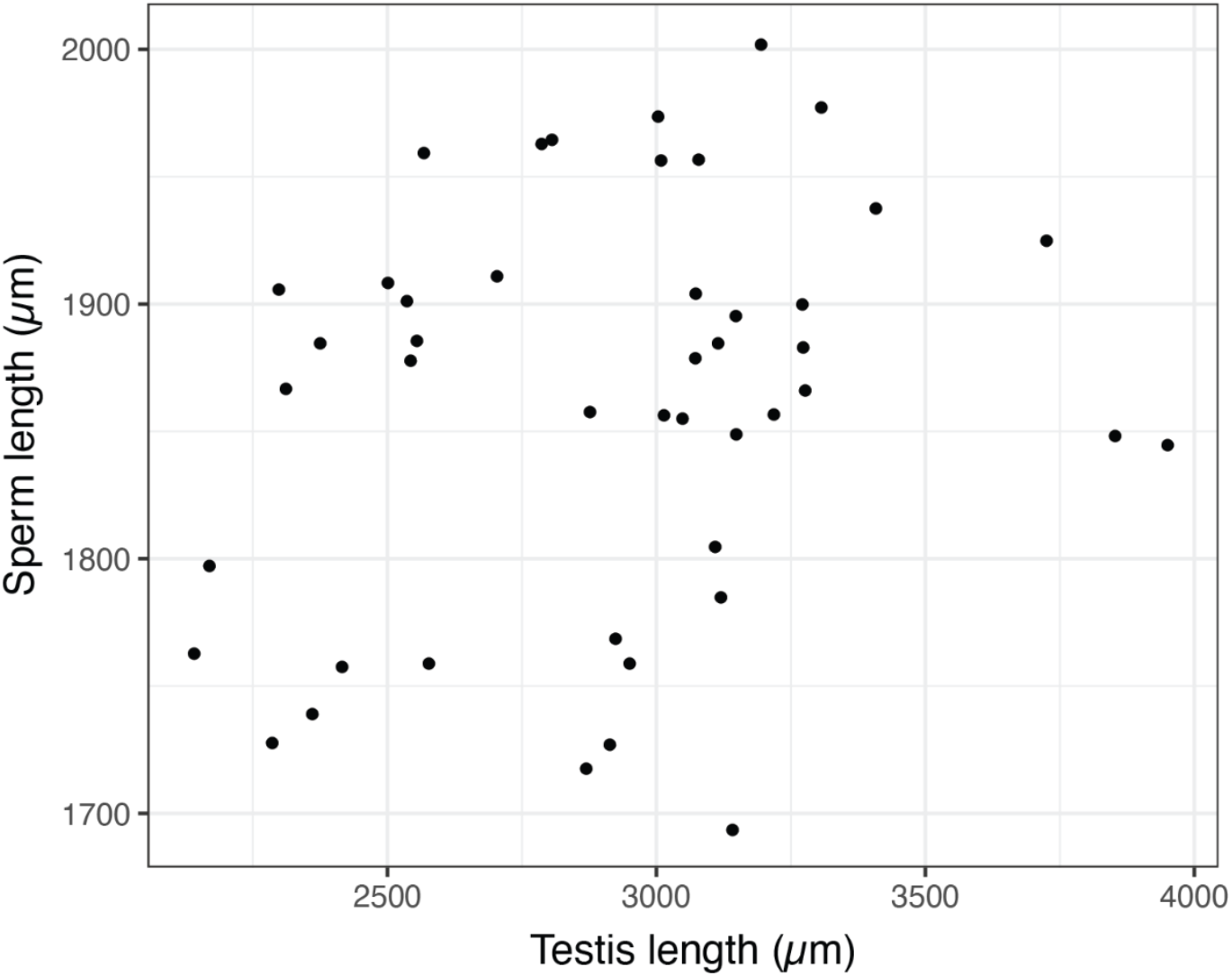
Sperm length is not correlated with testis length. For 45 wild type males, sperm length and testis length are not significantly correlated (*P* = 0.1164), suggesting that differential gene expression is not also a function of testis size.

**Figure S5.**
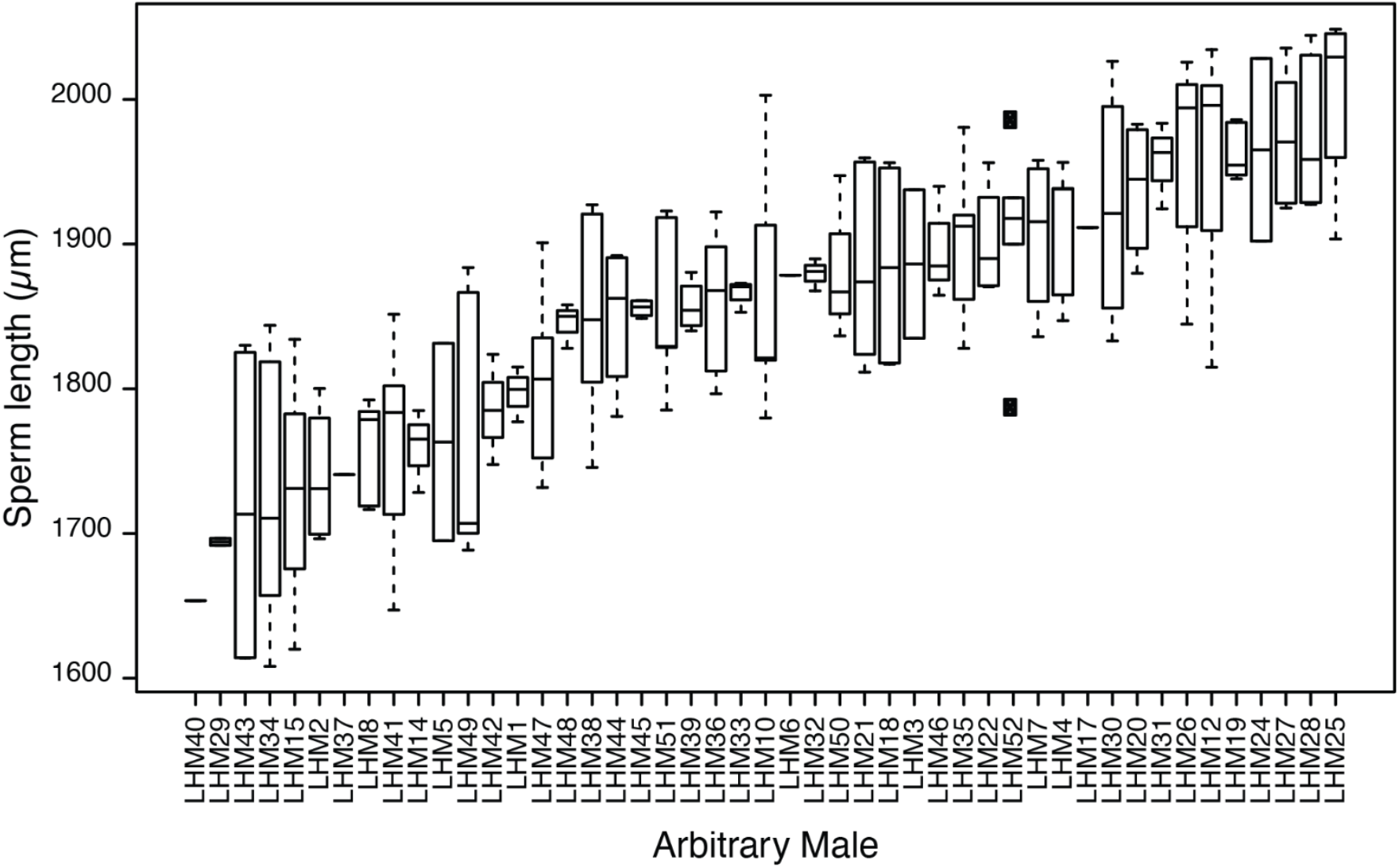
Distribution of sperm lengths within and among a subset of wild type males used to examine relationship between sperm length and testis length.

**Fig. S6.**
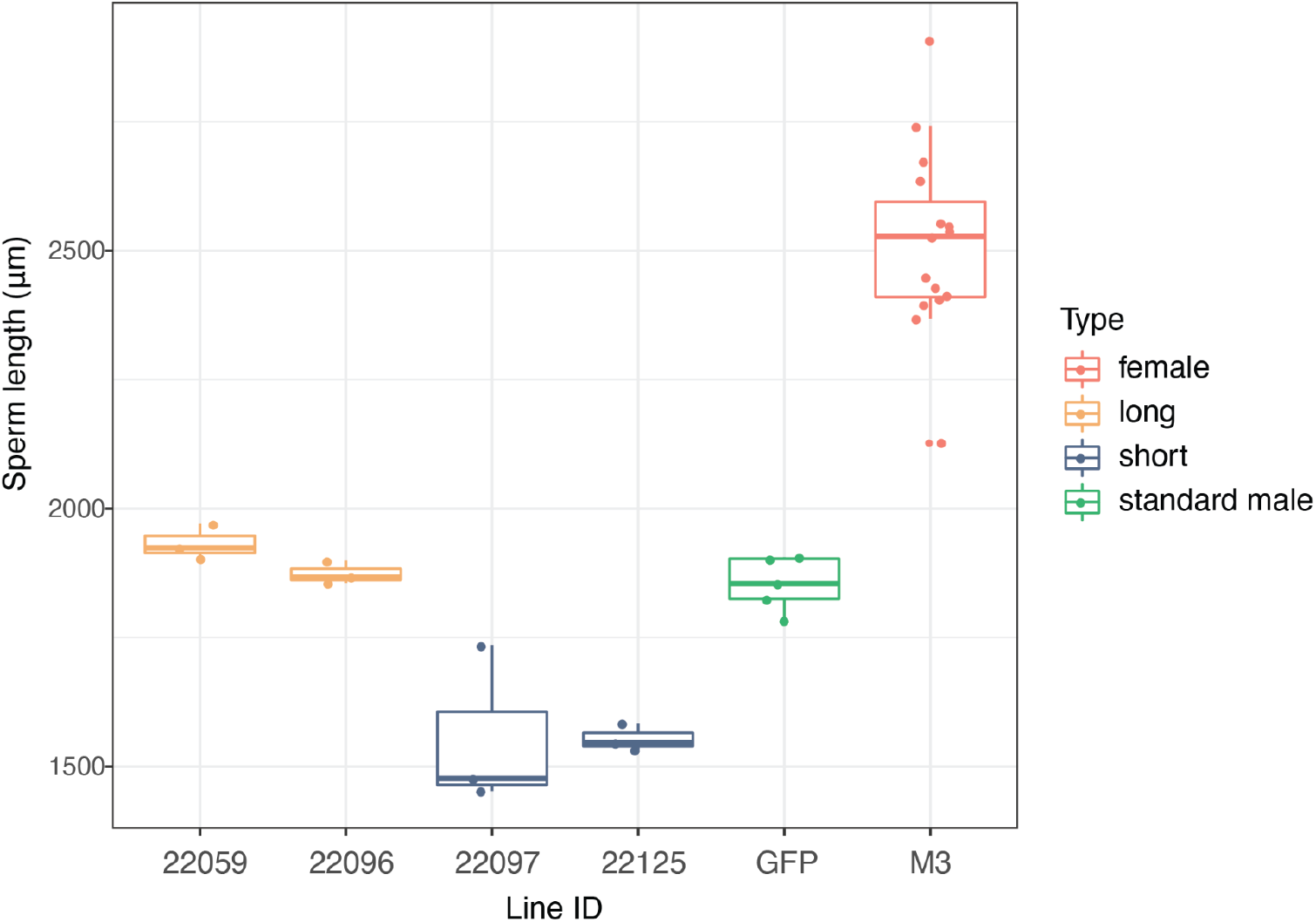
Sperm lengths for long sperm (22059, 22096) and short sperm DSPR RILs (22097, 22125), standard males (GFP), and SR lengths for standard females (M3).

**Table S1.** Summary of RNAseq libraries for *Drosophila* with short or long sperm producing testes. There were two replicate inbred isofemale lines for each sperm type: short (L08, L17) and long (H08, H20). Age indicates the days post-eclosion when males were dissected for RNA extraction.

| Sperm length | Sample | Male Age | RIN | Raw Reads | Mapped Reads |
| --- | --- | --- | --- | --- | --- |
| Short | L08_A | 6 days | 8.2 | 51,529,690 | 41,383,983 |
|  | L08_B | 6 days | 8.0 | 55,011,804 | 39,047,004 |
|  | L17_B | 4-6 days | 8.0 | 48,480,536 | 38,547,711 |
|  | L17_C | 5-6 days | 7.6 | 64,392,800 | 44,188,295 |
| Long | H08_A | 5-6 days | 7.7 | 68,174,383 | 51,032,772 |
|  | H08_B | 5-6 days | 7.3 | 58,918,527 | 41,370,628 |
|  | H08_C | 5-6 days | 6.3 | 64,118,639 | 49,661,976 |
|  | H20_A | 5-6 days | 7.9 | 55,229,124 | 37,227,203 |
|  | H20_B | 6 days | 8.3 | 56,032,571 | 45,009,281 |
|  | H20_C | 6 days | 8.3 | 51369699 | 39182439 |
| Total |  |  |  | 573,257,773 | 426,651,292 |

**Table S2.**
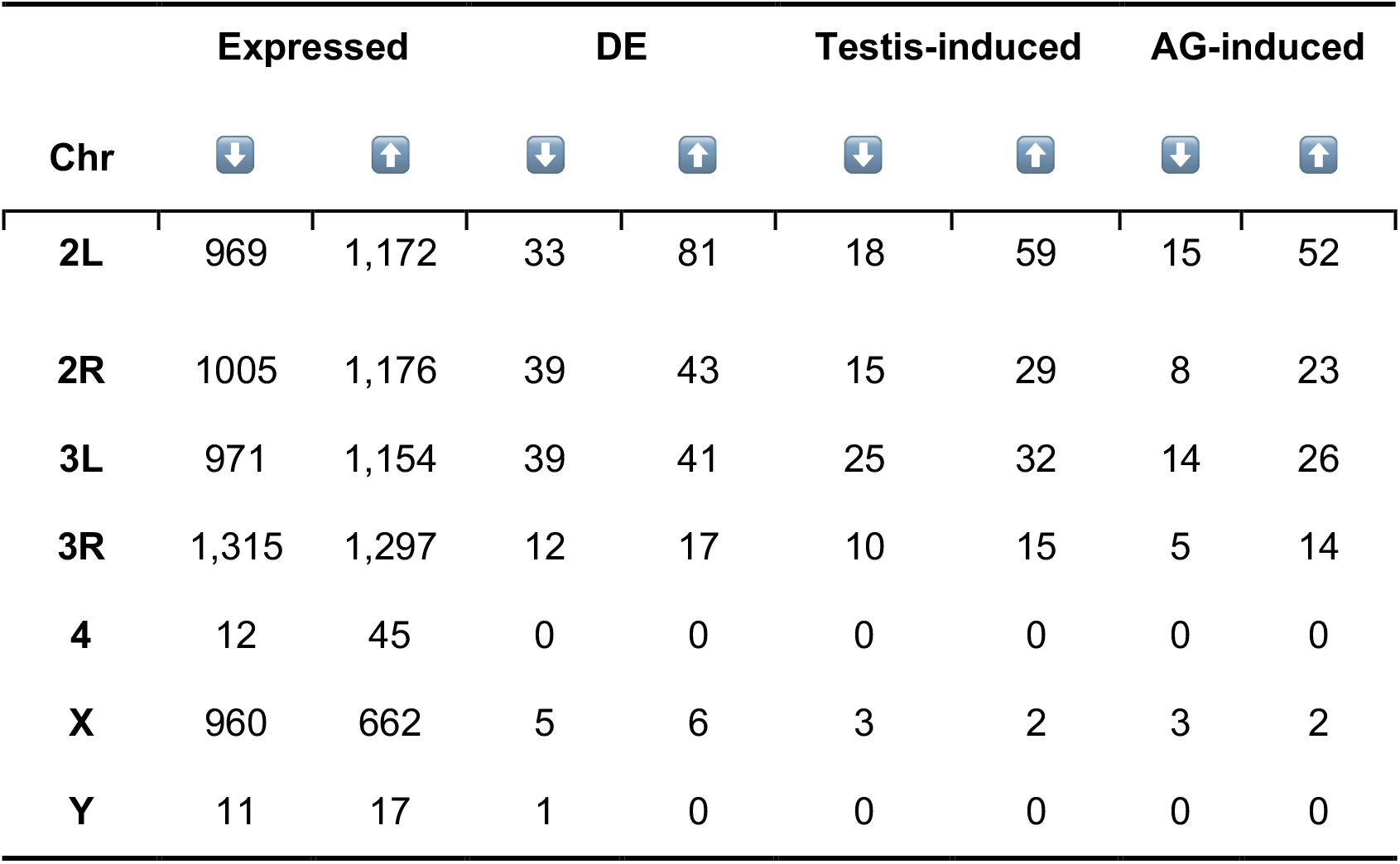
Summary of expression by chromosome. Genes that are expressed have an FPKM > 1 in a minimum of 4 samples. Genes that are induced in the testes (testis-induced) or accessory glands (AG-induced) had expression in that tissue that was higher than median expression in other tissues, based RNAseq tissue expression data from FlyBase. Arrows represent genes in each category that have positive (⬆) or negative (⬇) logFC in comparisons between short and long sperm producing testes. Positive logFC indicates higher expression in long sperm producing testes.

**Table S3.** Stocks used in this study.

| Stock(s) | Experiment | Provided by | Reference |
| --- | --- | --- | --- |
| L08, L17, H08, H20 | Testis transcriptomes | Co-author Manier | Zajitschek et al. (2019) |
| *Δ130/TM3, Sb, ry* | SP knockout | Mariana Wolfner | Liu & Kubli (2003) |
| *Δ325/TM3, Sb, ry* | SP knockout | Mariana Wolfner | Liu & Kubli (2003) |
| *Acp26Aa1* | Acp26Aa knockout | Mariana Wolfner | Herndon & Wolfner (1995) |
| *Df(2L)Exel6014* | Acp26Aa knockout | Mariana Wolfner | Parks et al. (2004) |
| *prd-GAL4* | AG knockout | Mariana Wolfner | Xue & Noll (2002) |
| *UAS-Rh1G69D* | AG knockout | Mariana Wolfner | Ryoo et al. (2007) |
| LHm | Testis size and sperm length | Co-author Manier | Rice et al. (2002) |
| 22059, 22096, 22097, 22125 | PMR | Stuart MacDonald | King et al. (2012) |
| M3 | PMR | Co-author Manier | This study |
| Canton-S GFP | PMR | Geoff Findlay | Chebbo et al. 2020 |

